# Representation Learning Methods for Single-Cell Microscopy are Confounded by Background Cells

**DOI:** 10.1101/2025.06.26.661577

**Authors:** Arushi Gupta, Alan Moses, Alex X. Lu

**Author notes:** Work primarily conducted during an internship at Microsoft Research New England (Arushi Gupta).

## Abstract

Deep learning models are widely used to extract feature representations from microscopy images. While these models are used for single-cell analyses, such as studying single-cell heterogeneity, they typically operate on image crops centered on individual cells with background information present, such as other cells, and it remains unclear to what extent the conclusions of single-cell analyses may be altered by this. In this paper, we introduce a novel evaluation framework that directly tests the robustness of crop-based models to background information. We create synthetic single-cell crops where the center cell’s localization is fixed and the background is swapped—e.g., with backgrounds from other protein localizations. We measure how different backgrounds affect localization classification performance using model-extracted features. Applying this framework to three leading models for single-cell microscopy for analyzing yeast protein localization, we find that all lack robustness to background cells. Localization classification accuracy drops by up to 15.8% when background cells differ in localization from the center cell compared to when the localization is the same. We further show that this lack of robustness can affect downstream biological analyses, such as the task of estimating proportions of cells for proteins with single-cell heterogeneity in localization. Ultimately, our framework provides a concrete way to evaluate single-cell model robustness to background information and highlights the importance of learning background-invariant features for reliable single-cell analysis.^1^

## 1 Introduction

Understanding single-cell variability can uncover important insights into underlying biological processes [1, 2, 3, 4, 5]. Single cells within the same population often differ in characteristics such as morphology, gene expression, and protein localization [1, 2, 3, 4, 5, 6]. In addition to having functional implications (e.g., it is a mechanism for populations to become robust to environmental changes [1, 2, 6]), single-cell variability allows researchers to observe asynchronous responses of cells (e.g., cells in different cell cycle stages), enabling them to identify regulatory processes without external intervention [3, 6]. As a result, single-cell variability has consistently been central to biological questions.

One key modality used to study single-cell variability is microscopy, as it can show the phenotype of many cells at single-cell resolution. To analyze microscopy images, researchers commonly quantify key features from cells in these images. In some cases, these features are computed from segmentations of individual cells, ensuring that the features reflect single cells in isolation [7]. More recently, many researchers have shifted toward using deep learning methods to automatically extract relevant features [8, 9, 10]. However, most deep learning approaches operate on single-cell crops, images that are centered on a single cell but include background information and possibly nearby cells. This raises the concern that these methods may inadvertently learn features from the surrounding context rather than the central single cell alone, potentially altering the conclusion of a single-cell analysis. In particular, this concern is amplified when a cell population is heterogeneous, because it raises the possibility that surrounding cells of different phenotypes may be influencing the interpretation of the centered cell.

To evaluate this concern, we propose a novel evaluation framework that detects if crop-based representation learning methods extract biologically robust single-cell representations. Our framework is based on the principle that a single cell’s representation should remain consistent despite changes to its background. Specifically, we swap single cells into images with different backgrounds and measure how much the resulting representations change. To demonstrate our framework, we apply it to models developed to understand single-cell protein localization using fluorescent microscopy because it is cell-specific and so it provides a strong setting for assessing whether learned representations depend solely on the individual cell.

We applied this framework to three deep learning-based single-cell feature representation methods developed to study protein localization in yeast cells: PIFiA [9], Paired Cell Inpainting [8], and DeepLoc [10]. We found that all three models rely significantly on background context. Specifically, across all models, classifiers trained on features extracted from background-swapped images consistently performed worse (by approximately 10-15%) than those trained on features extracted from unaltered images. We conduct rigorous controls that demonstrate these drops in performance are due to sensitivity to background cells with differing localization, as opposed to other explanations (e.g. artifacts from our swapping procedure, or batch effects).

Overall, our results demonstrate that existing deep learning models using single-cell crops are influenced by background context. We propose that our evaluation framework provides a useful standard for assessing and improving the robustness of future models in single-cell analysis tasks.

## 2 Related Work

### 2.1 Robustness in Single-Cell Microscopy Models

Prior work on the robustness of deep learning methods for single-cell microscopy has primarily focused on mitigating batch effects and image-level corruptions. Batch effects refer to systematic variations introduced by experimental conditions, such as differences in imaging instruments, staining protocols, or plate handling, that can obscure true biological differences [11, 12, 13]. Several studies have proposed methods to correct for batch effects, including semi-supervised learning strategies [11], batch-specific normalization techniques [12], and adaptations of data integration methods originally developed for single-cell transcriptomics [13]. Along with evaluating robustness to batch effects, several studies have benchmarked microscopy models against synthetic image degradations such as noise, blur, and optical aberrations [14, 15], designed to mimic common artifacts encountered in fluorescence microscopy imaging.

To our knowledge, no prior work has investigated robustness to background information in single-cell tasks. Instead, several studies have shown that incorporating background context can improve single-cell model performance when that context is biologically informative. For example, Snijder et al. [16] demonstrated that features of a single cell’s microenvironment, such as neighboring cell density and size, can be used to predict virus infection and endocytosis within that cell. Toth et al. [17] proposed a fisheye-style image transformation method that emphasizes pixels near the target cell, finding that it improved classification accuracy across several single-cell phenotyping tasks. In these studies, background context is causally or correlatively linked to the desired phenotype to predict. However, this relationship does not always hold. For example, the localization of cell cycle–regulated proteins is primarily determined by cell-intrinsic morphology (which reflects cell cycle stage) rather than surrounding context. Thus, when studying single-cell variability in settings like protein localization, using background information can introduce confounding factors that negatively impact performance, which we investigate.

### 2.2 Background Reliance in Natural Image Classification

While not explored in single-cell image settings, the impact of background information on the robustness of deep learning models is well-documented in natural image classification. Prior work has shown that deep learning models trained to classify objects often rely heavily on background features, and removing the background can significantly reduce classification accuracy [18, 19]. Xiao et al. [18] further demonstrated that such models can achieve non-trivial accuracy using background alone, and that adversarially modified backgrounds can induce misclassification even when the foreground object is unchanged. These findings highlight the risk that models trained on natural images may learn spurious correlations tied to background features.

Building on these findings, recent work has developed methods to measure and mitigate background reliance in natural image classification models. Moayeri et al. [20] introduced Relative Foreground Sensitivity, a metric that quantifies how much a model’s predictions rely on foreground versus background content. Their method uses saliency maps and ground truth segmentation masks to assess how closely model attention aligns with the object of interest. Bassi et al. [21] proposed a similar saliency-based approach that involves penalizing model attention to background regions to minimize background reliance during model training.

In our work, we measure the impact of background context in single-cell microscopy images by evaluating how model performance changes when the background information is either masked out or adversarially altered (similar to Xiao et al. [18]). This allows us to directly and simply test how background context affects model results, including in worst-case scenarios.

## 3 Methods

### 3.1 Dataset

We used the PIFiA dataset [9], which contains high-throughput images from the global yeast ORF-GFP collection, covering 4,049 unique strains. Each strain expresses a GFP-tagged protein along with fluorescent markers for the nucleus and cytoplasm. Cell images were collected from two biological replicates, each with four fields of view per GFP strain. We used data from the first replicate for all experiments.

We generated single-cell crops by segmenting cells in the cytoplasm channel to identify center coordinates, then extracting 64×64 pixel crops centered on each coordinate from the protein (GFP) channel. To support synthetic image generation (e.g., background swaps and masking), we applied the cytoplasm channel segmentation masks onto the GFP-channel crops. Segmentation was performed using YeastSpotter [22].

We applied two filtering steps to remove low-quality data for experiments. First, following [9], we excluded any crops with low protein signal by removing those whose mean GFP intensity fell below the 5th percentile of the global GFP intensity distribution, computed across all full-field GFP images. Second, we discarded segmented crops in which the mask covered less than 5% or more than 95% of the crop area, as these likely reflect failed segmentations (e.g., missing or oversized cell regions).

### 3.2 Image Generation

To assess the robustness of each single-cell representation model to background cell information, we generated synthetic single-cell image data where the center cell was from a different crop than the background cells.

Specifically, for every pair of localization categories (*l*_1_, *l*_2_), we did the following:

1. We randomly chose a protein *p*_1_ that localizes to *l*_1_ and a protein *p*_2_ that localizes to *l*_2_. When *l*_1_ = *l*_2_, we ensured that *p*_1_*≠ p*_2_.
2. We created 1,000 synthetic images. To create a single synthetic image, we randomly selected two single-cell GFP images, *s*_1_ and *s*_2_, from *p*_1_ and *p*_2_ respectively. We segmented the center cell *c*_1_ from *s*_1_, and superimposed it onto *s*_2_, which served as the background (we define the pixels not masked by *c*_1_ to be the background *b*_2_). To ensure consistent intensity scaling between *c*_1_ and *b*_2_), we standardize these pixels using the mean and standard deviation of pixel intensities calculated across all single-cell images for their respective proteins, *p*_1_ and *p*_2_.

Example synthetic background-swapped images are shown in Appendix C. For our localization categories, we use manual labels for the yeast ORF-GFP library by Huh et al. [23]; we filter these to just proteins with a single localization category (for 15 localization categories, as listed in Appendix B).

### 3.3 Assessing Robustness to Background Information

To evaluate the extent to which background context influences single-cell representations, we designed five experiments. In each, we trained linear classifiers to distinguish between features extracted from image crops with two different center cell localization classes (*c*_1_, *c*_2_), while systematically varying the background. We reasoned that evaluating accuracy of these classifiers would measure if salient features were being altered to the extent that they influenced classification boundaries. The specifics of the image crops used for each experiment type are below (we define an *aligned* background localization as a background where the cells share the same localization as the center cell):

- **Baseline:** Real single-cell crops with naturally aligning background localizations.
- **Same Localization Swap:** Synthetic single-cell crops where segmented center cells were placed on a single-cell crop from aligning background localizations *b*_1_ = *c*_1_, *b*_2_ = *c*_2_.
- **Different Localization Swap:** Synthetic single-cell crops where segmented center cells were placed on a single-cell crop from a third localization distinct from either localization classified by the linear classifier *b≠ c*_1_, *c*_2_.
- **Cell-Free Different Localization Swap:** Synthetic single-cell crops where segmented center cells were placed on background-only crops (no cells present) from a third localization *b≠ c*_1_, *c*_2_.
- **Masked Background:** Real single-cell crops with background pixels set to 0 using segmentation masks.

An example set of crops associated with all experiment types is shown in Figure 1A.

**Figure 1.**
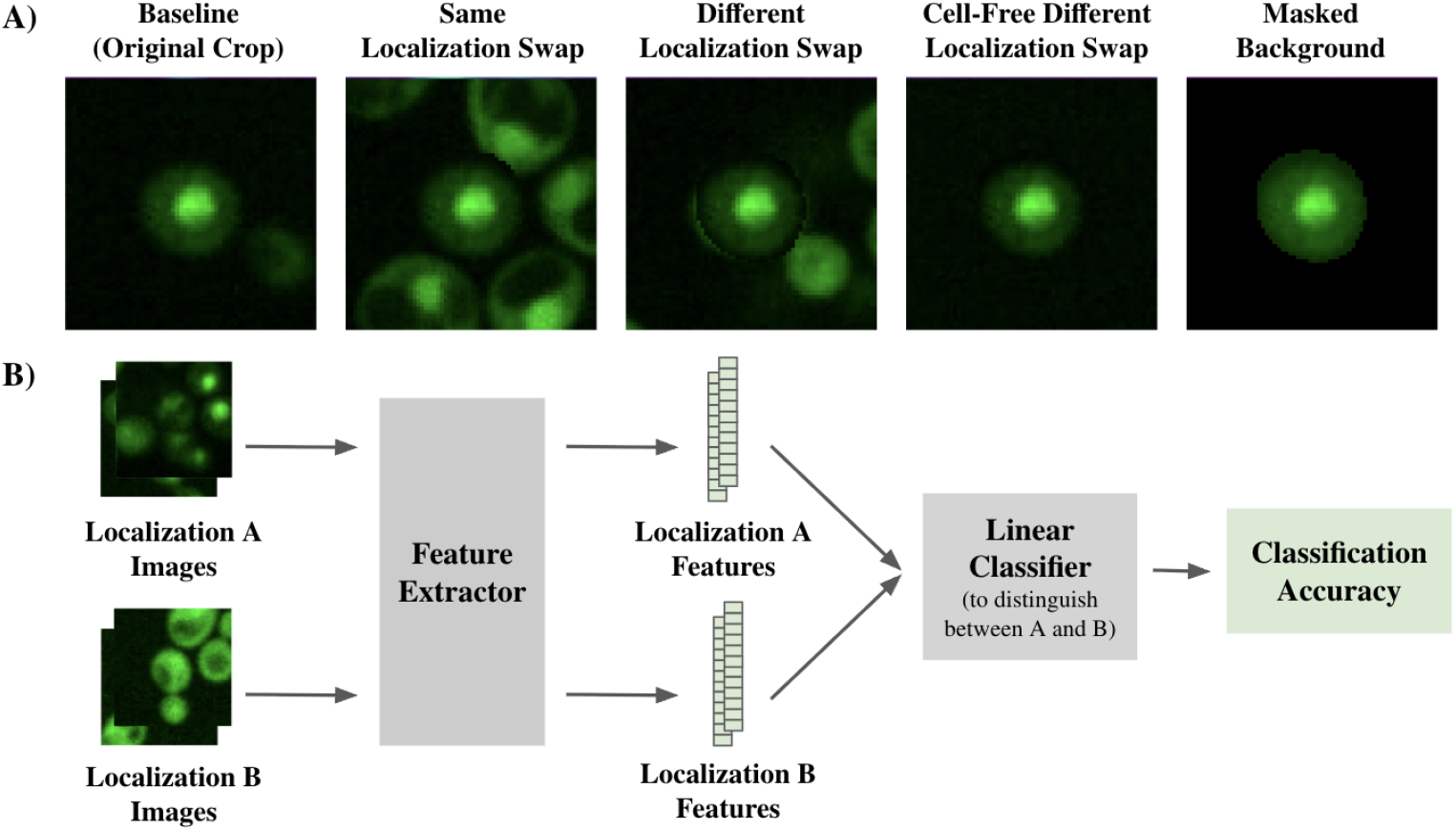
(A) We used 5 image settings to assess the role of background in single-cell representation models: unaltered crops, crops with swapped backgrounds from the same or different localization classes, crops with cell-free backgrounds from different localization classes, and crops with masked backgrounds. We show one example of these settings (images are colored in green to represent GFP, and rescaled to use the full intensity range). (B) In our evaluation, features were extracted from images of two different center cell localization classes and used to train a linear classifier. Accuracy was compared across image settings to assess the influence of background information on model representations.

### 3.4 Binary Classification

For each experiment type, we trained a separate Support Vector Machine (SVM) classifier with a linear kernel for every pair of center cell localizations (*c*_1_, *c*_2_), using data specific to that experiment type. In the Different Localization Swap and Cell-Free Different Localization Swap experiments, classifiers were trained for all triplets (*c*_1_, *c*_2_, *b*), where *b* denotes a background localization that differs from both *c*_1_ and *c*_2_.

For every pair or triplet, we constructed a dataset of 2,000 images, 1,000 with center cells from localization *c*_1_ and 1,000 from *c*_2_, with backgrounds defined by the experiment type. Each classifier was evaluated using 5-fold stratified cross-validation. Accuracy was computed for each fold and averaged across the five folds to obtain a final accuracy score for the classifier. To report overall results for each experiment type, we averaged the classification accuracies across all evaluated pairs or triplets. To report localization-level results, we averaged accuracies over all pairs or triplets in which the localization appeared as either *c*_1_ or *c*_2_.

For the Cell-Free Different Localization Swap experiment, since the goal was to check for the impact of batch effects (i.e. differences in imaging conditions) on model features, we additionally evaluated on only one 80/20 split of data, where we ensured that the set of proteins used in training set was different than those used from test set to isolate the potential impact of protein-level batch effects. The results are reported in the Appendix D.

### 3.5 Model Evaluation

We evaluated the feature representations of the PIFiA, Paired Cell Inpainting (PCI), and DeepLoc models for robustness to background information using our framework. For each model, we used average-pooled features from the intermediate layer that yielded the highest classification accuracy in classifying localizations. Full layer-wise accuracy comparisons are provided in Appendix A.

### 3.6 Multinomal Classification

In addition to our binary classifiers, which are used for our own evaluations, we trained multinomial classifiers to reproduce previous classifiers used to estimate proportions of cells for multi-localized proteins. Following Razdaibiedina et al. [9], we used data from proteins with a single subcellular localization, as manually annotated in Huh et al. [23]. Proteins labeled as “ambiguous” or assigned to localization classes with fewer than five proteins were excluded. From the selected 1,531 proteins, we extracted all valid single-cell crops and their corresponding masked versions, applying the filtering criteria described earlier (Section 3.1). Each crop was then passed through the PIFiA model to generate a 64-dimensional single-cell feature profile. This yielded two matched datasets, one from original crops and one from masked crops, each containing 2,668,343 feature profiles. We trained multinomial logistic regression models with the Adam optimizer, a learning rate of × 10^*−*3^, and a cross-entropy loss. Training was performed for 5 epochs.

## 4 Results

### 4.1 Same localization swaps do not degrade classification accuracy

Swapping the background of an image can introduce imaging artifacts, such as instances where the center cell overlaps with cells in the new background. To ensure that performance drops were not due to these artifacts (as opposed to surrounding cells where the phenotype of cells differed from the center cell), we compared the classification accuracy of same localization swaps to that of baseline images. We trained and evaluated three sets of binary classifiers, one set for each of the deep learning feature representations (PIFiA, Paired Cell Inpainting, and DeepLoc) we benchmark, using baseline images and *same localization swap images* (synthetic images where the background for center cells is swapped with that of another crop of the same localization).

We find that for the PIFiA and PCI feature representations, accuracy differences between the baseline test dataset and the same localization swap test dataset were minimal (less than 0.02) (Table 1, Figure 2A; full classification results are reported in Appendix B). For the DeepLoc feature representation, same localization swaps slightly improved classification accuracy (0.746 ± 0.030 versus 0.713 ± 0.029). Together, these results confirm that artifacts from background swapping do not adversely affect classification performance, suggesting that drops in classification accuracy can be attributed to changes in background content (as opposed to stretching representations and/or classification models out-of-distribution).

**Table 1:** Average localization classification accuracy (± 3 standard errors) per model across the 5 different experiment settings. Accuracy is averaged and standard error is calculated per localization class across all binary classification tasks involving that class.

| Model | Baseline | Diff Loc Swap | Same Loc Swap | Masked Background | Cell-Free Diff Loc Swap |
| --- | --- | --- | --- | --- | --- |
| PIFiA | 0.946±0.020 | 0.843±0.022 | 0.939±0.013 | 0.883±0.023 | 0.902±0.017 |
| PCI | 0.916±0.025 | 0.772±0.026 | 0.930±0.016 | 0.869±0.029 | 0.876±0.019 |
| DeepLoc | 0.713±0.029 | 0.612±0.025 | 0.746±0.030 | 0.677±0.029 | 0.689±0.023 |

**Figure 2.**
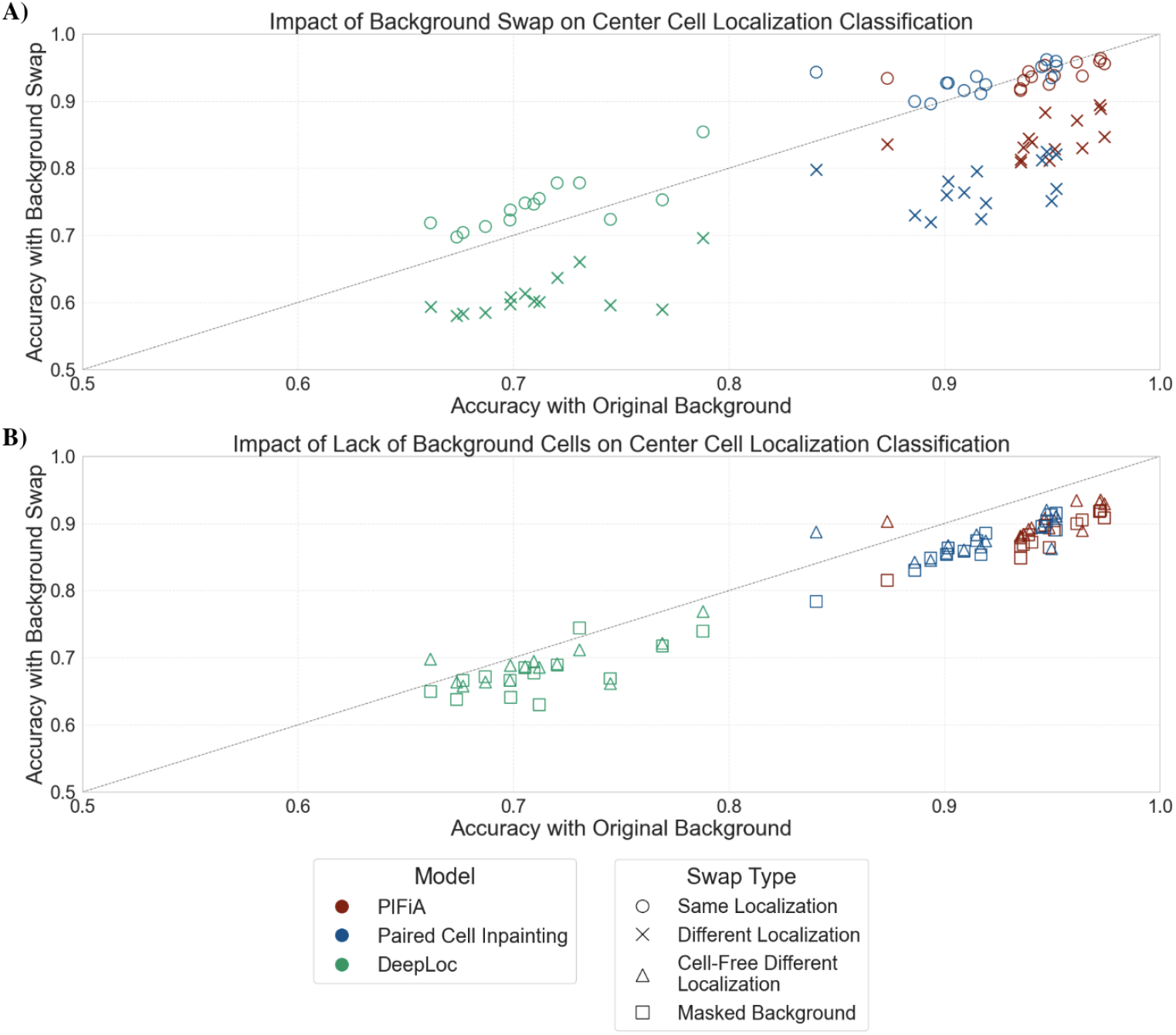
Each point shows the average classification accuracy for a specific localization class, calculated across all binary classification tasks involving that class. The x-axis shows accuracy using crops with their original backgrounds; the y-axis shows accuracy using crops with modified backgrounds. Points below the diagonal (x = y) indicate reduced accuracy with background changes. (A): Classifier performance when swapping cells into backgrounds with same versus different localization classes. (B): Classifier performance when swapping cells into backgrounds without cells or masking backgrounds.

### 4.2 Different localization swaps degrade classification accuracy

Having confirmed that artifacts from background swapping do not adversely affect model features, we next examined whether models are influenced by the localization of surrounding (background) cells. To do this, we next evaluated the classifier’s ability to generalize to *different localization swaps*. In this set-up, we evaluate if classifiers can distinguish between center cells with localizations *c*_1_ and *c*_2_ placed onto a background taken from a crop belonging to a third localization class *b* (with *b* ≠ *c*_1_, *c*_2_), intended to serve as a distractor phenotype. This experiment effectively simulates a single cell analysis in a heterogeneous setting, where the center cell of interest is surrounded by other, irrelevant cells of a different phenotype. As shown in Figure 2A, introducing different background localizations consistently and significantly reduced classification accuracy across all models and localizations. On average across localizations, accuracy dropped by 0.096 for PIFiA, 0.157 for PCI, and 0.134 for DeepLoc relative to the same localization swap setting (Table 1).

### 4.3 Decreases in classification accuracy are primarily explained by cells, not background

Next, we sought to establish that the drops in classification accuracy seen with different localization swaps were due to background cells of differing phenotypes, not because of batch effects. Deep representation learning methods can overfit to technical imaging differences, such as lighting, background intensity, or staining artifacts, that could be captured in model features. If single-cell crops of the same localization class share technical conditions, this could explain why accuracy drops in the different localization swap set-up, but not the same localization swap set-up. Indeed, we found some evidence that certain protein localization classes were more represented on some plates than others in the PIFiA dataset, suggesting that localization classes may share batch effects (Appendix D).

To rule this out, we next assessed classification accuracy under *cell-free different localization swaps*. In this set-up, center cells with localizations *c*_1_ and *c*_2_ were placed on background-only crops (i.e., no visible cells) from a third localization *b* (*b* ≠ *c*_1_, *c*_2_). These crops preserve any imaging artifacts associated with localization *b*, while removing the influence of background cells. As shown in Table 1, performance in the cell-free different localization swap setting was consistently higher than the different localization swap setting, suggesting that the performance drop can be largely attributed to surrounding cells of different localization, and not placing cells on a different background (that may reflect differing batch effects).

### 4.4 Surrounding cells improve classification accuracy when they are of the same localization

Having established that all models are sensitive to background cells, we reasoned that this may also have a positive impact on performance where background cells share the same phenotype as the center cell. To test this, we produced *masked background* crops, where we set all background pixels in each single-cell crop were set to zero, leaving only the center cell visible. Classification accuracy in this setting drops compared to the same localization swap setting across all models–on average, performance decreased by 0.05 for PIFiA, 0.06 for PCI, and 0.07 for DeepLoc (Table 1, Figure 2).

## 4.5 Masking surrounding cells yields differing estimates of proportions of phenotypes in multi-localized proteins

Through our evaluation framework, we showed that existing single-cell representation models are sensitive to background cells of differing phenotypes. We next sought to demonstrate how that may impact the conclusions of single cell analyses. To do this, we focused on understanding proteins that exhibit single-cell heterogeneity in localization, about 15% of yeast proteins [9]. Estimating the proportion of cells with localization to each compartment can shed biological insights (for example, some transcription factors can be heterogeneously localized, and the proportion of cells with a nuclear localization points to the strength of the response), so yeast protein databases [9, 24] will document the proportion of cells each compartment as predicted by classifiers on single cell microscopy crops [24, 10, 9].

Reproducing the procedures of Razdaibiedina et al. [9], we trained multinomial logistic regression classifiers for 15 localization classes on the PIFiA features (which typically performed the best in our previous analyses). The classifier is used to assign localization labels to individual cells, and then aggregated to estimate the proportion of cells in each compartment, for different proteins. Importantly, we train and predict using two classifiers: one on baseline image crops, and one on masked background image crops. This allows us to compare how predicted localization distributions shift when background context is removed.

We trained these classifiers on single-localized proteins. We then predicted the proportions of cells for 1,531 proteins annotated to be multiply localized by Huh et al. [23]. To quantify how much localization predictions differ when using baseline versus masked crops, we computed the Kullback–Leibler (KL) divergence between the predicted localization distributions for both models across all proteins. A higher KL value indicates a greater difference in predicted localization proportions. Across the 1,531 proteins, the KL divergence ranged from 0.003 to 9.75, with a median of 0.48 and a mean of 0.80. The full distribution of KL divergences is shown in Figure 3A. To contextualize these values, we calculated a baseline KL divergence between the original-crops model predictions and a uniform distribution over localization classes. For 223 out of 1,531 proteins (14.6%), the KL divergence between the two models exceeded this baseline, indicating that the models produced substantially different localization distribution predictions.

**Figure 3.**
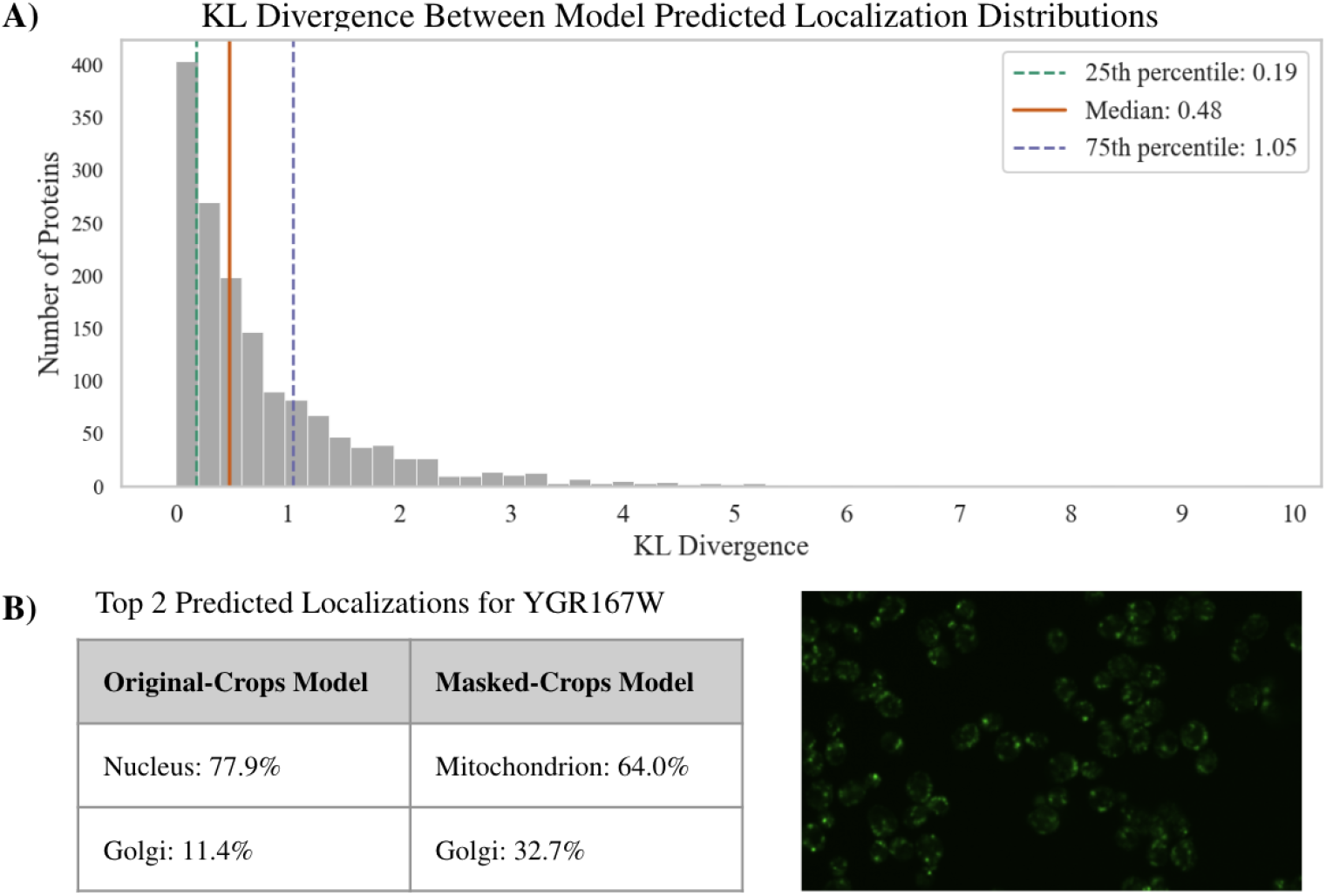
(A) Distribution of KL divergences calculated between protein localization distributions predicted by the original-crops and masked-crops logistic regression models. (B) Predictions of the original-crops and masked-crops models for YGR167W, and a representative cropped GFP image for this protein (right).

Finally, to better understand the nature of the disagreement between the two models, we manually examined the ten proteins with the largest KL divergence in predicted localization proportions. For each protein, we reviewed example images and compared the predicted distributions from the original-crops and masked-crops models. In 5 out of 10 cases, a blinded annotator rated the masked-crops model’s predictions as being more consistent with the visual appearance of the distribution of cells in full images, suggesting that the original-crops model’s outputs were distorted by background cells from differing localizations. An example is shown in Figure 3B.

These results reinforce our broader finding: background cells can significantly alter a single-cell model’s representation and predictions, and in some cases, removing background information yields more accurate estimates of localization proportions.

## 5 Discussion

Robust single-cell analysis is essential for understanding cellular variability and the biological processes that drive it. Across three leading feature extractors for studying protein localization in single yeast cells—PIFiA [9], Paired Cell Inpainting [8], and DeepLoc [10]—we consistently find that the learned representations lack robustness to background cells. Specifically, classification accuracy drops when background cells are from a different localization, and improves when they are from the same.

These findings have direct implications for past single-cell microscopy analyses. Analyses performed in prior works to evaluate localization classification frequently oversample single-localization proteins (as they are more abundant than heterogeneously localized proteins in yeast) [24, 10, 9, 8]. Our work demonstrates a bias where single-localization crops are classified more accurately than crops where surrounding cells are of different localizations, suggesting that these prior analyses may have overestimated our ability to classify single-cell protein localization. We show that classifier over-reliance on background can distort the conclusions of single-cell analyses: by comparing localization distributions for mixed-localization proteins using baseline versus masked single-cell crops, we show there are substantial differences for 15% of the proteins.

Our findings are particularly relevant given the emerging popularity of pooled screens, in which hundreds of proteins are expressed together within a single population of cells, enabling high-throughput, parallel analysis of protein localization under shared conditions [25]. As a result, microscopy images contain single-cell variability that reflects not just differences in localization of a single protein, but also that each cell may express completely different proteins. In this setting, accurate assignment of cells to protein identity depends on model features being robust to background information [25].

Ultimately, our study reveals a critical vulnerability in current representation learning methods for single-cell microscopy: a lack of robustness to background cells. We introduce a simple framework to quantify background robustness and benchmark three models. As single-cell imaging scales in complexity and throughput, ensuring that model features are robust to background cells for accurate downstream analysis is essential. Future work should consider the importance of isolating single cells where analyses operate on each cell independently.

## 6 Acknowledgements

We thank Brenda Andrews and Anastasia Razdaibiedina for their support and assistance with the data. We also thank David Alvarez-Melis, Enoch Luk, and the BioML team at Microsoft Research for helpful discussions and feedback.

## A Layer-wise Accuracy Comparisons

To determine which feature layer to use for each model in our background-swapping experiments, we evaluated the localization classification performance of all available layers from the PIFiA, Paired Cell Inpainting, and DeepLoc models. Results are shown in Supplementary Table 1.

**Supplementary Table 1.** Localization classification accuracy across feature layers for PIFiA, Paired Cell Inpainting (PCI), and DeepLoc. The best-performing layer for each model is shown in bold. Layers marked with “–” were not available for that model.

| Model | conv1 | conv2 | conv3 | conv4 | conv5 | conv6 | conv7 | conv8 | final |
| --- | --- | --- | --- | --- | --- | --- | --- | --- | --- |
| PIFiA | 0.9288 | 0.9571 | 0.9823 | 0.9870 | 0.9891 | 0.9879 | 0.9882 | **0.9896** | 0.9880 |
| PCI | 0.8227 | 0.9762 | 0.9828 | **0.9847** | 0.9832 | — | — | — | — |
| DeepLoc | — | — | 0.8100 | 0.8130 | **0.8436** | 0.8356 | 0.8296 | 0.8185 | 0.7891 |

### Evaluation setup

We constructed a balanced dataset across 15 protein localization categories. For each category, we selected three representative proteins that localized exclusively to that category, based on localization percentages reported by Razdaibiedina et al. [9], and applied a localization purity threshold. From each protein, we randomly sampled 1,000 single-cell images.

**Supplementary Table 2.**
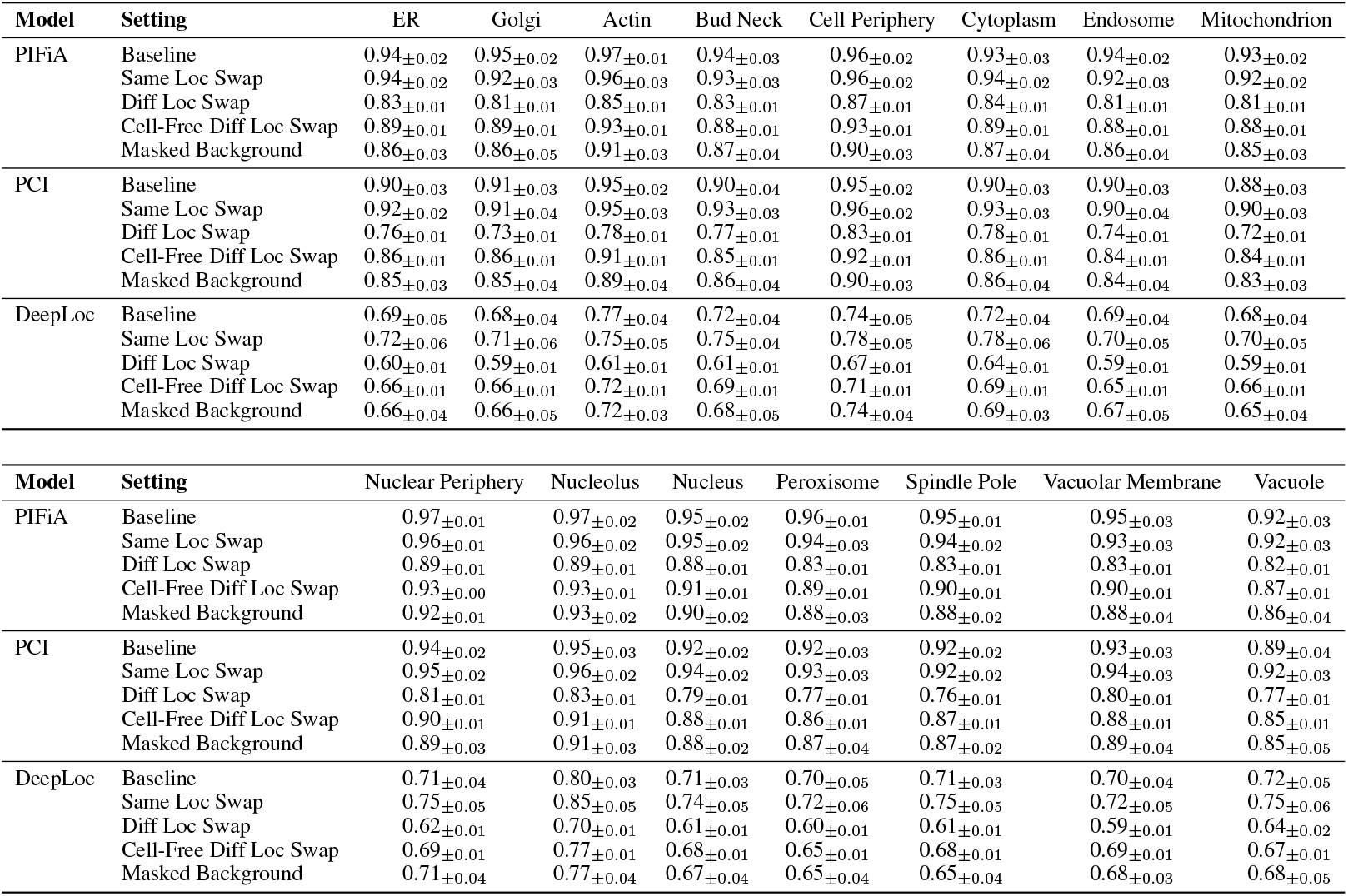
Localization-wise classification accuracy (± 3 standard errors) for each model across five experimental settings. Accuracy is averaged per localization class across all binary classification tasks involving that class.

| Model | Setting | ER | Golgi | Actin | Bud Neck | Cell Periphery | Cytoplasm | Endosome | Mitochondrion |
| --- | --- | --- | --- | --- | --- | --- | --- | --- | --- |
| PIFiA | Baseline | 0.94±0.02 | 0.95±0.02 | 0.97±0.01 | 0.94±0.03 | 0.96±0.02 | 0.93±0.03 | 0.94±0.02 | 0.93±0.02 |
| | Same Loc Swap | 0.94±0.02 | 0.92±0.03 | 0.96±0.03 | 0.93±0.03 | 0.96±0.02 | 0.94±0.02 | 0.92±0.03 | 0.92±0.02 |
| | Diff Loc Swap | 0.83±0.01 | 0.81±0.01 | 0.85±0.01 | 0.83±0.01 | 0.87±0.01 | 0.84±0.01 | 0.81±0.01 | 0.81±0.01 |
| | Cell-Free Diff Loc Swap | 0.89±0.01 | 0.89±0.01 | 0.93±0.01 | 0.88±0.01 | 0.93±0.01 | 0.89±0.01 | 0.88±0.01 | 0.88±0.01 |
| | Masked Background | 0.86±0.03 | 0.86±0.05 | 0.91±0.03 | 0.87±0.04 | 0.90±0.03 | 0.87±0.04 | 0.86±0.04 | 0.85±0.03 |
| PCI | Baseline | 0.90±0.03 | 0.91±0.03 | 0.95±0.02 | 0.90±0.04 | 0.95±0.02 | 0.90±0.03 | 0.90±0.03 | 0.88±0.03 |
| | Same Loc Swap | 0.92±0.02 | 0.91±0.04 | 0.95±0.03 | 0.93±0.03 | 0.96±0.02 | 0.93±0.03 | 0.90±0.04 | 0.90±0.03 |
| | Diff Loc Swap | 0.76±0.01 | 0.73±0.01 | 0.78±0.01 | 0.77±0.01 | 0.83±0.01 | 0.78±0.01 | 0.74±0.01 | 0.72±0.01 |
| | Cell-Free Diff Loc Swap | 0.86±0.01 | 0.86±0.01 | 0.91±0.01 | 0.85±0.01 | 0.92±0.01 | 0.86±0.01 | 0.84±0.01 | 0.84±0.01 |
| | Masked Background | 0.85±0.03 | 0.85±0.04 | 0.89±0.04 | 0.86±0.04 | 0.90±0.03 | 0.86±0.04 | 0.84±0.04 | 0.83±0.03 |
| DeepLoc | Baseline | 0.69±0.05 | 0.68±0.04 | 0.77±0.04 | 0.72±0.04 | 0.74±0.05 | 0.72±0.04 | 0.69±0.04 | 0.68±0.04 |
| | Same Loc Swap | 0.72±0.06 | 0.71±0.06 | 0.75±0.05 | 0.75±0.04 | 0.78±0.05 | 0.78±0.06 | 0.70±0.05 | 0.70±0.05 |
| | Diff Loc Swap | 0.60±0.01 | 0.59±0.01 | 0.61±0.01 | 0.61±0.01 | 0.67±0.01 | 0.64±0.01 | 0.59±0.01 | 0.59±0.01 |
| | Cell-Free Diff Loc Swap | 0.66±0.01 | 0.66±0.01 | 0.72±0.01 | 0.69±0.01 | 0.71±0.01 | 0.69±0.01 | 0.65±0.01 | 0.66±0.01 |
| | Masked Background | 0.66±0.04 | 0.66±0.05 | 0.72±0.03 | 0.68±0.05 | 0.74±0.04 | 0.69±0.03 | 0.67±0.05 | 0.65±0.04 |

| Model | Setting | Nuclear Periphery | Nucleolus | Nucleus | Peroxisome | Spindle Pole | Vacuolar Membrane | Vacuole |
| --- | --- | --- | --- | --- | --- | --- | --- | --- |
| PIFiA | Baseline | 0.97±0.01 | 0.97±0.02 | 0.95±0.02 | 0.96±0.01 | 0.95±0.01 | 0.95±0.03 | 0.92±0.03 |
| | Same Loc Swap | 0.96±0.01 | 0.96±0.02 | 0.95±0.02 | 0.94±0.03 | 0.94±0.02 | 0.93±0.03 | 0.92±0.03 |
| | Diff Loc Swap | 0.89±0.01 | 0.89±0.01 | 0.88±0.01 | 0.83±0.01 | 0.83±0.01 | 0.83±0.01 | 0.82±0.01 |
| | Cell-Free Diff Loc Swap | 0.93±0.00 | 0.93±0.01 | 0.91±0.01 | 0.89±0.01 | 0.90±0.01 | 0.90±0.01 | 0.87±0.01 |
| | Masked Background | 0.92±0.01 | 0.93±0.02 | 0.90±0.02 | 0.88±0.03 | 0.88±0.02 | 0.88±0.04 | 0.86±0.04 |
| PCI | Baseline | 0.94±0.02 | 0.95±0.03 | 0.92±0.02 | 0.92±0.03 | 0.92±0.02 | 0.93±0.03 | 0.89±0.04 |
| | Same Loc Swap | 0.95±0.02 | 0.96±0.02 | 0.94±0.02 | 0.93±0.03 | 0.92±0.02 | 0.94±0.03 | 0.92±0.03 |
| | Diff Loc Swap | 0.81±0.01 | 0.83±0.01 | 0.79±0.01 | 0.77±0.01 | 0.76±0.01 | 0.80±0.01 | 0.77±0.01 |
| | Cell-Free Diff Loc Swap | 0.90±0.01 | 0.91±0.01 | 0.88±0.01 | 0.86±0.01 | 0.87±0.01 | 0.88±0.01 | 0.85±0.01 |
| | Masked Background | 0.89±0.03 | 0.91±0.03 | 0.88±0.02 | 0.87±0.04 | 0.87±0.02 | 0.89±0.04 | 0.85±0.05 |
| DeepLoc | Baseline | 0.71±0.04 | 0.80±0.03 | 0.71±0.03 | 0.70±0.05 | 0.71±0.03 | 0.70±0.04 | 0.72±0.05 |
| | Same Loc Swap | 0.75±0.05 | 0.85±0.05 | 0.74±0.05 | 0.72±0.06 | 0.75±0.05 | 0.72±0.05 | 0.75±0.06 |
| | Diff Loc Swap | 0.62±0.01 | 0.70±0.01 | 0.61±0.01 | 0.60±0.01 | 0.61±0.01 | 0.59±0.01 | 0.64±0.02 |
| | Cell-Free Diff Loc Swap | 0.69±0.01 | 0.77±0.01 | 0.68±0.01 | 0.65±0.01 | 0.68±0.01 | 0.69±0.01 | 0.67±0.01 |
| | Masked Background | 0.71±0.04 | 0.77±0.04 | 0.67±0.04 | 0.65±0.04 | 0.65±0.04 | 0.68±0.03 | 0.68±0.05 |

### Classification task

For each model, we extracted features from each available layer and trained a linear SVM to classify between all pairs of localization categories. Each pairwise classifier was trained on a balanced set of 2,000 images (1,000 per class). Performance was evaluated using 5-fold stratified cross-validation. Accuracy was averaged across folds and across all localization pairs. All features from convolutional layers were average-pooled across spatial dimensions.

### Model Specifics

For Paired Cell Inpainting, which follows an encoder–decoder architecture, we evaluated only the intermediate convolutional layers of the encoder, as these were used for feature extraction at inference time in the original paper. For PIFiA and DeepLoc, along with evaluating intermediate layers, we additionally evaluated the final feature representation from the second fully connected (FC) layer, which precedes the classification head. In PIFiA, this corresponds to the 64-dimensional feature profile used in downstream analyses. In DeepLoc, this is the 512-dimensional output from the second of three FC layers.

## B Localization-Wise Classification Accuracies Across All Experiments

In Supplementary Table 2, we report full localization-wise classification accuracies for each model across all five experimental settings: Baseline, Same Localization Swap, Different Localization Swap, Masked Background, and Cell-Free Different Localization Swap.

## C Visualizations

In Supplementary Figure 1, we display randomly sampled synthetic single-cell crops from the Different Localization Swap dataset to show the visual variability and potential imaging artifacts introduced during background swapping.

Supplementary Figure 2 shows examples of misclassified single-cell crops from the Cell-Free Different Localization Swap experiment, where classifiers were tasked with distinguishing between two center cell localizations, with backgrounds containing cells from a third, different localization.

These examples reveal several contributing factors to misclassification. First, we see examples where the center cell is clearly consistent with its labeled localization, supporting the idea that the presence of background cells with a different localization is responsible for the misclassification. Second, we observe examples where the center cell itself is visually ambiguous. For instance, in row 1, column 5, the center cell localization is labeled as cytoplasm but appears as though it could be nucleus, which it is incorrectly classified as. Third, we observe imaging artifacts introduced during the background swap process, which reduce the visual clarity of the center cell, possibly contributing to the misclassification. For example, in row 3, column 3, the center cell appears visually unclear as there is a background cell partially behind it.

The second and third factors also apply to the data in the Same Localization Swap experiment. Thus, the accuracy drop observed in Different Localization Swap relative to Same Localization Swap can be primarily attributed to the first factor: the presence of background cells with a different localization.

**Supplementary Figure 1.**
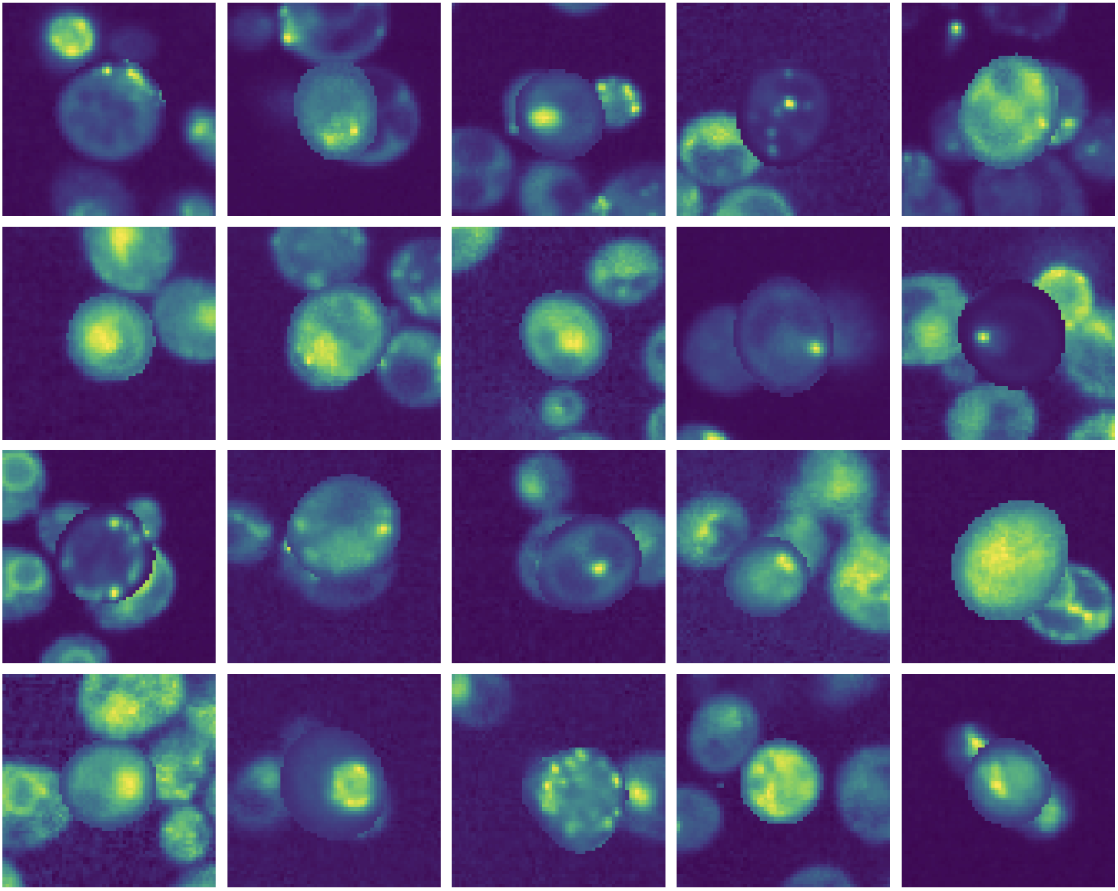
Randomly sampled synthetic single-cell crops from the Different Localization Swap dataset. Each image shows a segmented center cell from one localization imposed on a single-cell crop from another localization.

**Supplementary Figure 2.**
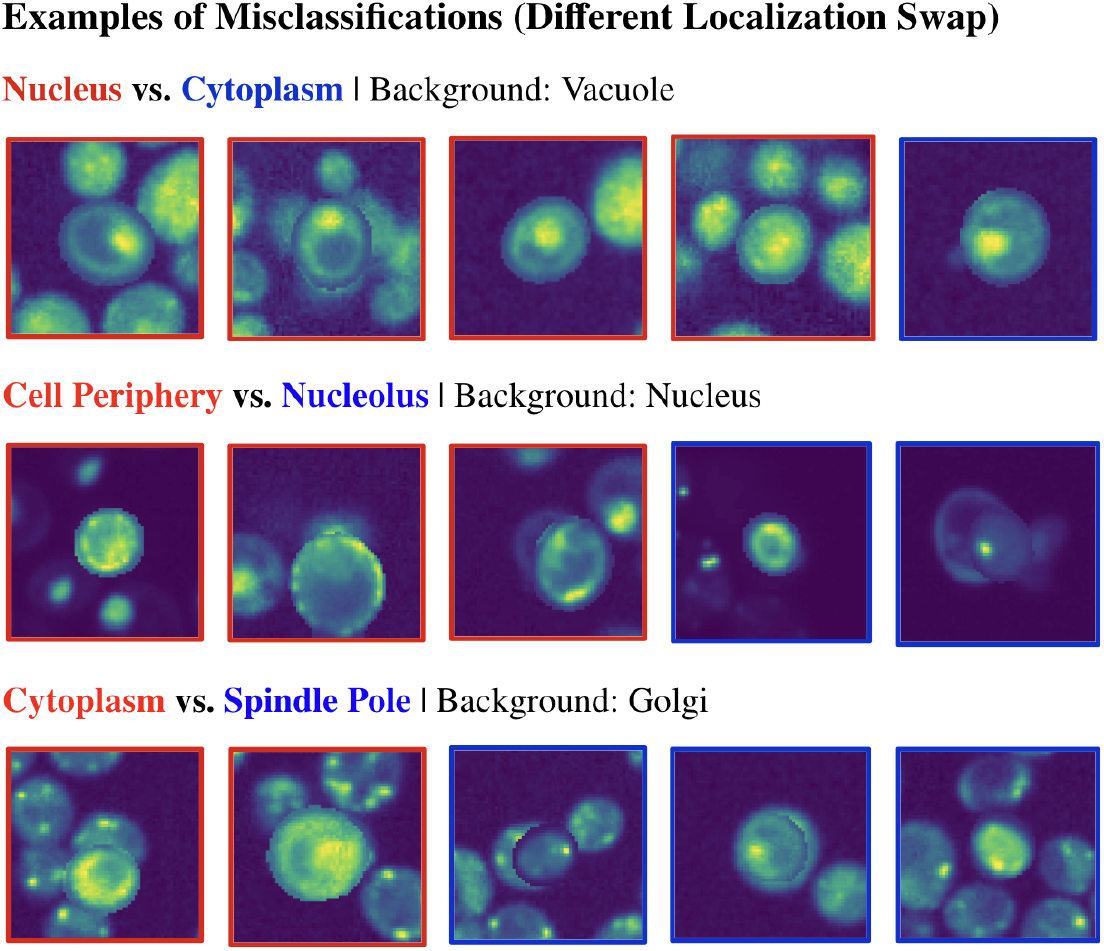
Examples of misclassified single-cell crops in the Different Localization Swap experiment. All shown examples are misclassifications, and the border color indicates the ground truth center cell localization: red for the first listed class and blue for the second.

## D Cell-Free Different Localization Swap: Additional Information

In this section, we provide additional information and analysis for the Cell-Free Different Localization Swap experiment.

### D.1 Data Generation

To construct the Cell-Free Different Localization Swap dataset, we extracted background-only crops (i.e., regions without visible cells) from the full-field GFP images of each protein in the PIFiA dataset. We excluded image borders to avoid partial or poorly segmented cells, and then identified candidate crops using a fixed stride of 32 pixels. To ensure these crops were cell-free, we discarded any that were within 50 pixels of a detected cell center and further filtered out low-quality regions using a pixel intensity variance threshold.

From the resulting background crops, we generated the final dataset using a two-step sampling procedure: we first randomly selected a protein with a known localization to the desired class, and then sampled one of its background crops. If no valid crop was available for the chosen protein, we repeated the sampling.

### D.2 Analysis of Plates and Localizations

To investigate potential localization-specific batch effects in our dataset, we analyzed how single-localization proteins are distributed across imaging plates. Supplementary Figure 3 displays a heatmap of localization proportions, normalized within each localization category (i.e., each row sums to one), showing how frequently proteins from a given localization were imaged on each plate.

We observe substantial skew in these distributions: several localization classes—such as bud neck, cell periphery, nuclear periphery, and peroxisome—are heavily concentrated on specific plates. For example, over 50% of cell periphery-labeled proteins appear on plate 10, and over 40% of bud neck-labeled proteins appear on plate 1.

This plate-level clustering suggests that proteins of the same localization were often imaged under shared experimental conditions. As a result, single-cell crops of these proteins may exhibit confounding signals tied to imaging artifacts rather than biology. To isolate and assess the impact of such effects, we introduced the Cell-Free Different Localization Swap experiment, where segmented cells are imposed onto cell-free crops from a different localization.

### D.3 Protein-Heldout Evaluation

In the main paper, we reported results for the Cell-Free Different Localization Swap from 5-fold cross-validation, where the training and test sets may contain cells from the same protein. However, it’s possible that model features capture protein-level batch effects.

To more rigorously test for the presence of batch effects, we additionally evaluated model performance using a protein-held-out 80/20 split. In this setup, the training and test sets contain disjoint sets of proteins, ensuring that any protein-specific imaging conditions cannot be exploited during classification.

Supplementary Table 3 displays the localization-wise classification accuracies for both settings. Results are similar for both, suggesting that models are likely not relying on protein-level batch effects.

**Supplementary Figure 3.**
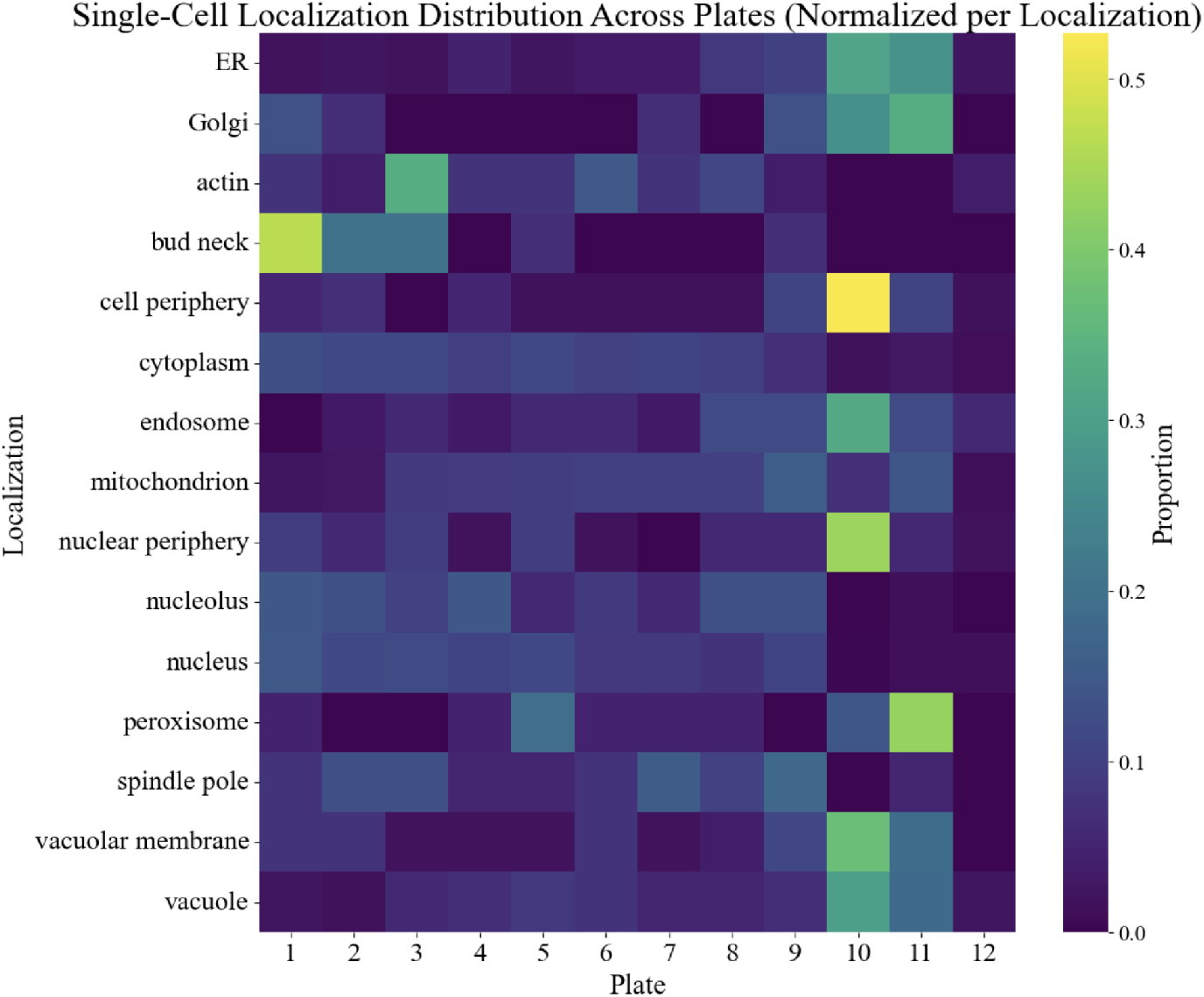
Heatmap showing the distribution of single-localization proteins across imaging plates. Values are normalized per localization row, such that each row (localization) sums to 1. Brighter colors indicate a higher proportion of that localization on a given plate.

**Supplementary Table 3.** Localization-wise classification accuracy for the Cell-Free Different Localization Swap experiment. Rows show the accuracies from a protein-held-out 80/20 split (no protein overlap) and 5-fold cross-validation (with potential protein overlap).

| Model | Evaluation Type | ER | Golgi | Actin | Bud Neck | Cell Periphery | Cytoplasm | Endosome | Mitochondrion |
| --- | --- | --- | --- | --- | --- | --- | --- | --- | --- |
| PIFiA | Protein-Held-Out Split | 0.873 | 0.816 | 0.890 | 0.880 | 0.877 | 0.878 | 0.843 | 0.864 |
|  | 5-Fold Cross-Validation | 0.890 | 0.891 | 0.929 | 0.883 | 0.930 | 0.887 | 0.877 | 0.879 |
| PCI | Protein-Held-Out Split | 0.839 | 0.765 | 0.870 | 0.853 | 0.874 | 0.850 | 0.818 | 0.824 |
|  | 5-Fold Cross-Validation | 0.858 | 0.861 | 0.906 | 0.852 | 0.920 | 0.861 | 0.843 | 0.840 |
| DeepLoc | Protein-Held-Out Split | 0.680 | 0.619 | 0.751 | 0.720 | 0.734 | 0.702 | 0.676 | 0.661 |
|  | 5-Fold Cross-Validation | 0.664 | 0.656 | 0.723 | 0.688 | 0.707 | 0.691 | 0.654 | 0.657 |

| Model | Evaluation Type | Nuclear Periphery | Nucleolus | Nucleus | Peroxisome | Spindle Pole | Vacuolar Membrane | Vacuole |
| --- | --- | --- | --- | --- | --- | --- | --- | --- |
| PIFiA | Protein-Held-Out Split | 0.932 | 0.915 | 0.908 | 0.945 | 0.888 | 0.823 | 0.852 |
|  | 5-Fold Cross-Validation | 0.934 | 0.934 | 0.909 | 0.893 | 0.899 | 0.898 | 0.869 |
| PCI | Protein-Held-Out Split | 0.909 | 0.895 | 0.874 | 0.916 | 0.853 | 0.758 | 0.825 |
|  | 5-Fold Cross-Validation | 0.896 | 0.911 | 0.883 | 0.864 | 0.870 | 0.879 | 0.849 |
| DeepLoc | Protein-Held-Out Split | 0.727 | 0.795 | 0.677 | 0.689 | 0.669 | 0.660 | 0.688 |
|  | 5-Fold Cross-Validation | 0.685 | 0.769 | 0.683 | 0.654 | 0.676 | 0.686 | 0.669 |

## E Multinomial Logistic Regression: Additional Results

### E.1 Performance on Held-Out Test Set

As described in Section 4.5, we trained multinomial logistic regression classifiers using single-cell features from proteins annotated as single-localization. While our main analysis focused on applying these classifiers to multiply localized proteins, we initially evaluated their performance by holding out 20% of the single-localization data as a test set.

Supplementary Table 4 reports the precision, recall, and F1-score for each localization class on this test set, comparing models trained on original image crops (which include background cells) versus masked crops (where background cells are removed). Consistent with the background swapping experiments, we observe that masking the background leads to a substantial drop in classification performance across most localizations. These results further validate our finding that the presence of background cells of the same localization improves localization classification.

**Supplementary Table 4.** Performance of logistic regression classifiers trained on original versus masked single-cell crops. Evaluation is conducted on a held-out test set of data of single-localization proteins. Metrics reported are per-class precision, recall, and F1-score.

| Localization | Original-Crop Model |  |  | Masked-Crop Model |  |  |
| --- | --- | --- | --- | --- | --- | --- |
|  | Precision | Recall | F1 | Precision | Recall | F1 |
| ER | 0.87 | 0.76 | 0.81 | 0.66 | 0.48 | 0.56 |
| Golgi | 0.78 | 0.58 | 0.66 | 0.38 | 0.03 | 0.05 |
| Actin | 0.89 | 0.82 | 0.85 | 0.69 | 0.28 | 0.40 |
| Bud neck | 0.80 | 0.30 | 0.44 | 0.70 | 0.04 | 0.07 |
| Cell periphery | 0.85 | 0.63 | 0.73 | 0.57 | 0.22 | 0.32 |
| Cytoplasm | 0.83 | 0.94 | 0.88 | 0.67 | 0.90 | 0.76 |
| Endosome | 0.77 | 0.55 | 0.65 | 0.67 | 0.05 | 0.10 |
| Mitochondrion | 0.88 | 0.84 | 0.86 | 0.64 | 0.65 | 0.65 |
| Nuclear periphery | 0.86 | 0.82 | 0.84 | 0.82 | 0.49 | 0.61 |
| Nucleolus | 0.91 | 0.85 | 0.88 | 0.75 | 0.64 | 0.69 |
| Nucleus | 0.90 | 0.90 | 0.90 | 0.83 | 0.79 | 0.81 |
| Peroxisome | 0.81 | 0.69 | 0.74 | 0.52 | 0.21 | 0.30 |
| Spindle pole | 0.81 | 0.69 | 0.75 | 0.64 | 0.33 | 0.43 |
| Vacuolar membrane | 0.84 | 0.65 | 0.74 | 0.67 | 0.28 | 0.40 |
| Vacuole | 0.70 | 0.59 | 0.64 | 0.47 | 0.19 | 0.27 |

## Footnotes

1 Our code is available at: https://github.com/microsoft/microscopy-sc-robustness

